# Epistasis among clustered lineage-specific amino acid substitutions in the Drosophila Trio protein

**DOI:** 10.1101/2025.09.29.679247

**Authors:** Flora Borne, Andrew M. Taverner, Peter Andolfatto

## Abstract

Intramolecular epistasis is increasingly recognized as a key factor shaping patterns of evolutionary rate variation among protein sites and constraining adaptive evolution. While genome-wide analyses have revealed that intramolecular epistatic interactions can drive the spatial clustering of amino acid substitutions, direct empirical evidence for such interactions and their evolutionary consequences remains limited. Using a population genetic screen for spatially-clustered and lineage-specific adaptive amino acid substitutions in *Drosophila* proteins, we systematically identify experimentally tractable candidates for functional analysis. As proof of concept, we focus on the Trio protein, a Rho guanine nucleotide exchange factor that exhibits three spatially-clustered putatively adaptive amino acid substitutions in the *D. melanogaster* lineage. By systematically reconstructing evolutionary intermediates *in vivo* using genome editing, we find that all possible intermediate states exhibit reduced viability and/or locomotor defects, providing strong evidence for epistatic constraints on evolutionary trajectories. Notably, these deleterious effects are recessive, suggesting that intermediate combinations of epistatically interacting amino acid substitutions can accumulate in heterozygotes prior to fixation, thereby circumventing apparent constraints imposed by maladaptive intermediate states. Together, these findings provide a rare empirical view of the fitness landscape shaped by intramolecular epistasis and establish a framework for investigating the constraints on adaptive protein evolution in diploid multicellular organisms.

**Author Summary:** Proteins fold into three-dimensional structures that are essential for their function. Because these structures depend on interactions among amino acids, the fitness effect of a mutation at one site can depend on the amino acid states at other sites. Such dependencies constrain the paths that protein evolution can take, whether evolution proceeds neutrally or adaptively. Although intramolecular epistasis has been demonstrated in microbial systems and *in vitro*, direct experimental evidence for such constraints in diploid multicellular organisms *in vivo* is rare. Here, we identify Drosophila proteins that exhibit clusters of closely-spaced putatively adaptive amino acid substitutions and focus experimental analyses on one example, Trio, a Rho guanine nucleotide exchange factor. Using genome editing to reconstruct evolutionary intermediates in the *D. melanogaster* lineage, we find that all intermediate versions of Trio reduce viability or impair locomotor function. Importantly, these harmful effects are recessive, suggesting that they could be masked when paired with an ancestral version of the protein. This implies that individual amino acid changes may persist in heterozygotes within populations and later combine to contribute to adaptation. If the recessivity of deleterious intermediates along adaptive evolutionary paths proves to be widespread, it could have important implications for our mechanistic understanding of adaptive protein evolution.

## Introduction

The rate of protein evolution varies substantially among species, among proteins and among sites within proteins. This rate variation has historically been interpreted in terms of “neutral”, or “nearly neutral” models of protein evolution, which depend on the proportion of newly arising amino acid altering mutations that are neutral with respect to protein function or effectively neutral due to genetic drift (1,2). While nearly neutral models can account for several important aspects of protein evolution (3), a debate raged for decades about the relative contribution of positive selection to protein evolution. The development and application of the McDonald-Kreitman test (4) and similar frameworks (5–7) suggested that protein divergence is well in excess of that predicted under neutral models of protein evolution in Drosophila and other species (5,8–13). While the McDonald-Kreitman and similar tests suffer from uncertainty of the population size over time (14,15), amino acid substitutions also exhibit genetic hitchhiking signatures at closely linked neutral sites, confirming an important role for positive selection in protein evolution (11,16–20).

Evolutionary rate variation within proteins is primarily believed to arise from differences in selective constraint, reflecting the need to maintain the integrity of specific protein domains and their functions (21–23). However, since positive selection also contributes substantially to protein divergence, this raises the possibility that clustered substitutions may sometimes result from repeated rounds of adaptive substitution in particular functional regions, or from the co-fixation of nearly neutral variants through genetic hitchhiking. Indeed, several studies have reported that positively selected amino acid substitutions tend to occur in close proximity within protein structures (24–29).

Most of the population genetic models assume that sites contribute independently to fitness. However, when considering the three-dimensional structure of proteins and their role in determining function, it becomes clear that amino acid residues act collectively to produce a coherent functional state. As sequences evolve, this coherence must be maintained or, when novel functions arise, re-established. Thus, residues are constrained to evolve non-independently, and the phenotypic and fitness effects of new amino acid–altering mutations depend on the history of substitutions at interacting sites. This dependence, termed intramolecular epistasis, is expected to play an important role in constraining protein evolutionary trajectories. This principle was first formalized in the covarion model, which proposed that only a subset of codons is free to vary at any given time and that this set shifts as substitutions occur (30–32). Since spatially proximate amino acids are more likely to directly interact, substitutions that compensate or permit one another are expected to co-occur closely in sequence space or atomic distance in folded proteins. Consistent with this prediction, recent studies have revealed that spatially clustered amino acid substitutions exhibit characteristic signatures of epistasis and compensatory evolution (33–38). Together, these findings suggest that intramolecular epistasis not only contributes to variation in evolutionary rate but also leaves a distinct spatial footprint in the form of clustered substitutions.

Experimental work has further confirmed the central role of intramolecular epistasis in protein evolution. A landmark study by Weinreich et al. (2006) showed that interactions among five mutations in β-lactamase dramatically constrained the number of traversable evolutionary paths to antibiotic resistance (39). Similarly, Ortlund et al. (2007) demonstrated that the glucocorticoid receptor evolved new hormone specificity in vertebrates via “stability-mediated” epistasis, in which nearly-neutral permissive mutations fixed stochastically and stabilized local structures, enabling subsequent adaptive substitutions to fix (40). Later studies using ancestral protein reconstruction, experimental evolution, and large-scale mutational scans have confirmed that epistatic interactions can strongly influence evolutionary trajectories (41–48). Furthermore, a few studies provide evidence of epistatic substitutions occurring in close spatial proximity, echoing the clustering patterns predicted from statistical analyses. For example, compensatory mutations have been observed near adaptive substitutions conferring toxin resistance in the poison frog nicotinic acetylcholine receptor (49), and similar patterns appear along the evolution of vertebrate proteins like p53, hemoglobin, lysozyme and Na⁺,K⁺-ATPase (50–54).

This experimental work has been instrumental in demonstrating the pervasiveness of epistasis and in revealing key principles of higher-order and global epistasis governing fitness landscapes. However, most of these studies have explored epistatic dynamics in the context of protein activity and stability *in vitro* (41–44,55,56), or microbial fitness *in vivo* (46–48). Consequently, a substantial gap still exists in our understanding of how interacting substitutions accumulate in diploid multicellular organisms where dominance and interactions across multiple phenotypic levels may also strongly influence evolutionary outcomes.

To our knowledge, Na⁺,K⁺-ATPase (NKA) resistance to cardiotonic steroids (CTS) in insects is the only case where epistasis in protein adaptation has been directly demonstrated *in vivo* in a multicellular organism (57–59). In CTS-adapted insects, adaptation frequently targets a highly conserved domain of the NKA alpha-subunit. The most common resistance-conferring substitutions—at positions 111 and 122—almost always occur after a permissive substitution at position 119 (A119S), which reduces the negative pleiotropic effects of these resistance mutations (57,58). Moreover, *in vivo* engineering of firefly CTS-resistance substitutions into Drosophila revealed similar deleterious pleiotropic effects, which were alleviated by additional background substitutions that occur within a span of 12 amino acid residues, but are not directly related to CTS resistance (59). Further, it was shown that the deleterious effects of CTS resistance-conferring substitutions appear to be recessive, potentially explaining how they might persist in populations long enough to allow secondary compensating substitutions to arise and enable their fixation. This work highlighted the use of *in vivo* models in revealing pleiotropic effects on higher order phenotypes that are more closely related to organismal fitness (e.g viability, fertility, and behavior) and would be missed in *in vitro* assays.

The case of CTS-resistant NKAs is a striking example of the role of epistasis in protein adaptation, but it remains unclear how broadly these findings apply to other proteins. To better understand how pleiotropy, epistasis, and dominance shape protein evolution during adaptation, new diploid *in vivo* models are needed. To this end, we systematically searched the *Drosophila melanogaster* genome for proteins that, like NKA, are highly conserved yet contain lineage-specific clusters of positively-selected amino acid substitutions. As a case study, we focus on one such cluster in the highly-conserved protein Trio, a Rho guanine nucleotide exchange factor involved in signalling pathways that regulate several important developmental processes. Using scarless CRISPR-Cas9 genome editing, we engineered Drosophila strains carrying all possible intermediate combinations of substitutions in this cluster, enabling us to dissect the epistatic and dominance interactions involved.

## Results

### Lineage-specific spatially-clustered amino acid substitutions in *D. melanogaster* proteins

To identify proteins with lineage-specific, spatially-clustered and putatively adaptive amino acid substitutions in *D. melanogaster*, we applied the Model-Averaged Site Selection with Poisson Random Field (MASS-PRF; (26)) to multiple sequence alignments for 9137 Drosophila proteins (See Methods). MASS-PRF estimates a site-by-site scaled selection coefficient (γ *= 2Ns*) based on polymorphism and divergence clustering models. While the McDonald-Kreitman test is typically applied to whole proteins, MASS-PRF is designed to quantify variation in intensities of selection across a protein sequence, highlighting potentially localized targets of adaptive protein evolution. We identified 919 proteins containing at least one cluster of putatively adaptive substitutions specific to the *D. melanogaster* lineage (Fig1, Fig S1, See Methods). The identification of these spatially-clustered amino acid substitutions provides the opportunity to investigate potential epistatic interactions among individual substitutions within such clusters. We estimated that 679 proteins contain experimentally tractable clusters consisting of 2 or 3 substitutions located less than 20 amino acids away from each other.

As proof of concept, we focus on a putatively adaptive cluster of amino acid substitutions in the protein Trio (Fig 1A) as a tractable example. Trio is a Rho-guanine nucleotide exchange factor (GEF) that regulates several signaling pathways involved in functions such as neuronal migration, axonal outgrowth, axon guidance, and synaptogenesis in *D. melanogaster* (60,61), *C. elegans* (62) and in humans (63,64). Missense and nonsense mutations in *trio* can cause neurodevelopmental diseases such as intellectual disability and autism spectrum disorders in humans (63). In *D. melanogaster*, *trio* mutants show specific neurodevelopmental disorders resulting in locomotion behavior, leg and wing formation defects but also more general phenotypes such as prepupal lethality and sterility (65,66).

**Figure 1.**
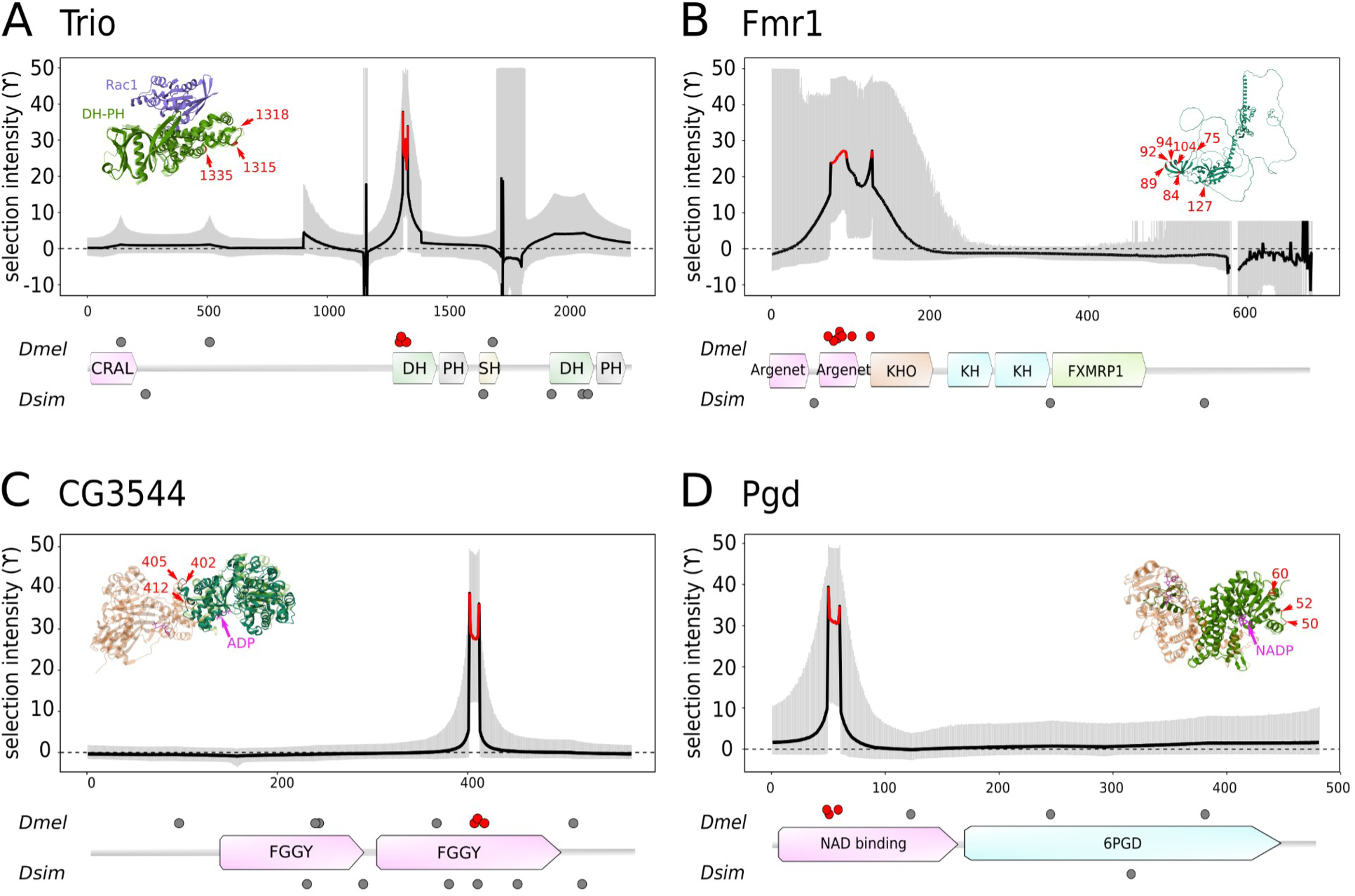
Examples of proteins exhibiting *D. melanogaster* lineage-specific clusters of putatively adaptive amino acid substitutions. Plotted are profiles of selection intensity (γ = *2Ns*) across four *D. melanogaster* proteins inferred with MASS-PRF (26). The black line corresponds to the model-averaged γ and the grey areas indicate 95% model uncertainty interval. Red lines indicate regions for which the 95% lower bound of γ > 4, which corresponds to a false positive rate of <0.1 (see Methods). Protein structure models are represented below each selection intensity plot. Functional domains are represented by colored boxes with their names in black. Lineage-specific amino acid substitutions are represented by dark gray dots and red for the cluster of interest (*D. melanogaster* above and *D. simulans* below, respectively). AlphaFold predictions of the 3D structure of the *D. melanogaster* proteins are inset. The prediction has been superimposed onto the structures of the homologous proteins (transparent) with ligands and in oligomerized form, when available (See enlarged structure Fig S2). The positions of the clustered amino acid substitutions are highlighted with red arrows. (A) Trio, a Rho guanine nucleotide exchange factor. The AlphaFold prediction of the first DH-PH domain of Trio is represented in green and is superimposed onto the human Trio DH-PH structure (transparent green) in complex with Rac1 (blue) (PDB 7SJ4, (67)). (B) Fragile X messenger ribonucleoprotein 1 (Fmr1). (C) CG3544, a xylulokinase. The AlphaFold prediction is represented in green and is superimposed on the xylulokinase structure of *Chromobacterium violaceum* (transparent orange and green) in complex with ADP (pink) (PDB 3KZB). (D) Phosphogluconate dehydrogenase (Pgd). The AlphaFold structure prediction is represented in green and is superimposed onto the dimerized structure of the human Pgd (transparent orange and green) in complex with NADP (pink) (PDB 2JKV).

The *D. melanogaster* and *D. simulans* Trio proteins differ by 11 amino acid substitutions, six of which occur along the *D. melanogaster* lineage (Fig 1A; Fig S3). A cluster of three amino acid substitutions (V1315A; N1318S; R1335K), exhibiting lineage-specific positive selection in *D. melanog*aster, is located in the first GTPase binding domain DH-PH of the protein. This domain is otherwise highly conserved and these three substitutions are the only amino acid changes observed across the *D. melanogaster-D. yakuba* species group, which shared a common ancestor ∼10 Mya (67–69) (Fig 1A; Fig S3). Further, these sites are monomorphic among 947 wild-derived *D. melanogaster* genome sequences from 30 populations worldwide (70), these three substitutions are monomorphic and we detected only three nonsynonymous singleton variants across the domain, indicating strong purifying selection.

While this cluster of amino acid substitutions are inferred to have been fixed by positive selection using the site-specific MASS-PRF approach, there is no additional evidence for recent or recurrent positive selection acting on the Trio protein. For example, application of the standard McDonald-Kreitman test and a branch-site model (implemented in PAML (71)) to the entire protein do not reject neutrality and there is no obvious signature of recent selective sweep (Fig S4). Further, the amino acid substitutions in the Trio cluster appear to be relatively conservative, preserving both polarity and charge properties (Fig S3), though there is a trend towards incorporation of smaller functional groups. Prediction tools assessing the impact of single substitutions on protein folding-stability were applied to the *D. melanogaster* AlphaFold-predicted structure of the DH-PH domain (72–74). These analyses did not identify any substitutions predicted to strongly destabilize the DH-PH domain, with all predicted ΔΔG values ranging between –1 and +1 kcal/mol (Table S1). We note, however, that these predictions rely on a modeled AlphaFold structure and should therefore be interpreted with caution.

### Strong epistasis among clustered amino acid substitutions in the Trio protein

To investigate potential phenotypic effects associated with the three amino acid substitutions in the Trio DH-PH domain (V1315A, N1318S and R1335K), we implemented scarless CRISPR-cas9/piggyBAC genome editing to the native Trio protein in *D. melanogaster* (see Methods). We edited *D. melanogaster* strains representing all possible combinations of this cluster of substitutions in the Trio protein from the ancestral state “VNR” to the current state “ASK” (Fig 2C). To avoid effects of genomic background, we homogenized the lines by crossing to well-characterized inbred strains (see Methods and Fig S5).

**Figure 2.**
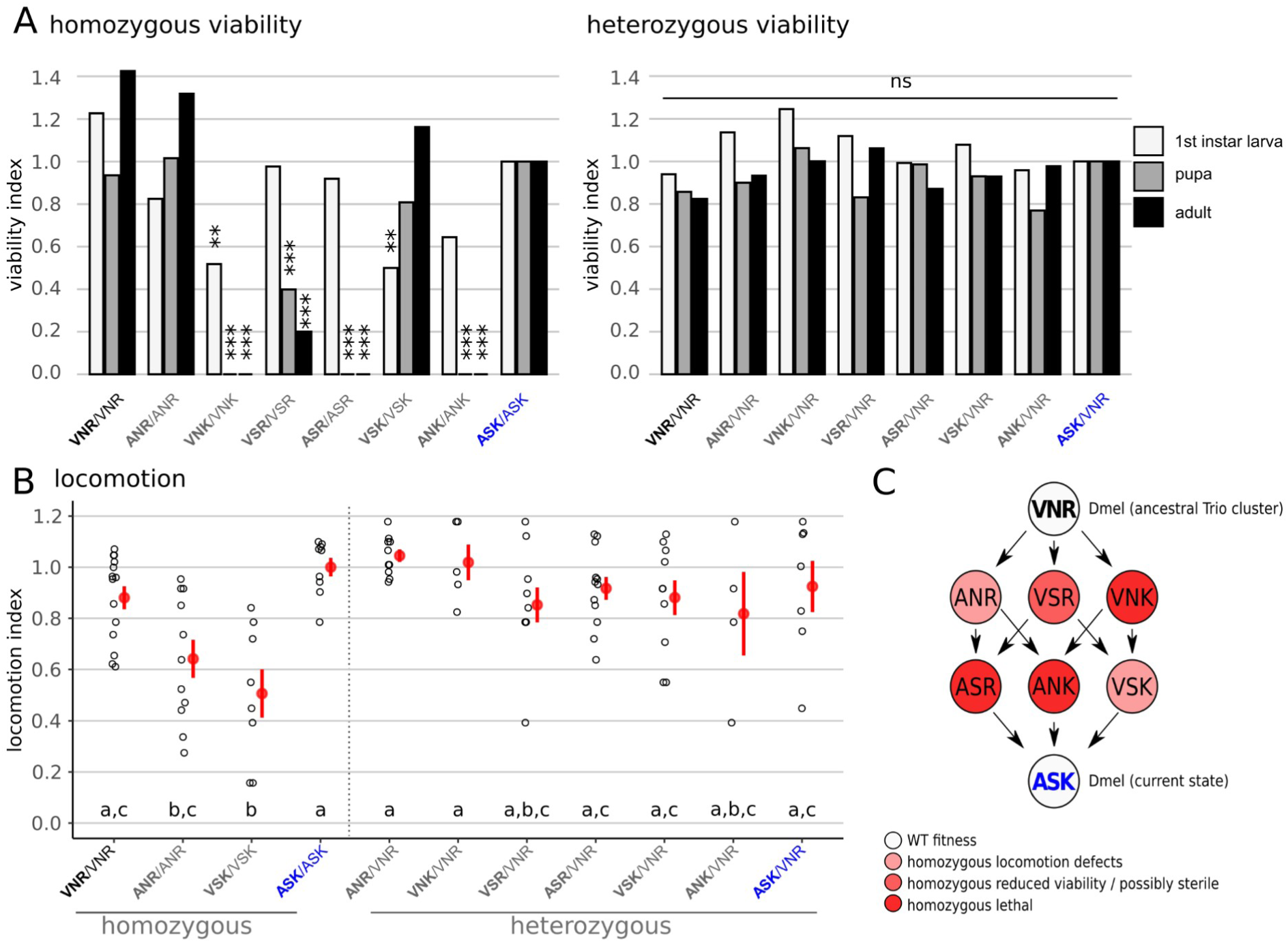
Evolutionary intermediates in the path from ancestral to extant *D. melanogaster* Trio sequences exhibit recessive deficits in fitness-related phenotypes. (A) Viability of homozygous and heterozygous first instar larvae, pupae and adults. Plotted is the proportion of YFP- individuals normalized relative to the ASK haplotype (see Methods). Asterisks indicate adjusted p-values determined by Fisher’s Exact tests comparing each genotype to the ASK haplotype (*** p<0.001; ** p<0.01; * p<0.05; p-values are Bonferroni-corrected). (B) Locomotion defects in homozygous and heterozygous individuals. A locomotion index (y-axis) is measured using a negative geotaxis assay and normalized relative to the ASK/ASK genotype (see Methods). Each dot represents a group of 5-10 males. The red dot and whiskers represent the mean and standard error, respectively. Two strains that share a letter are not significantly different from each other (Kruskall-Wallis followed by Dunn test, p>0.05). (C) Diagram representing all available genetic sequential substitution paths from the ancestral haplotype VNR to the extant *D. melanogaster* haplotype ASK. Colors of the circle represent the fitness of each haplotype based on viability and locomotion results.

To identify fitness effects associated with the three substitutions, and potential epistatic interactions among them, we first investigated homozygous effects of substitutions at these three sites on two fitness-related traits (viability and locomotion). Interestingly, most of the intermediate haplotypes showed severe defects in larval and adult viability. In particular, VNK, ASR and ANK flies are homozygous lethal and most individuals do not reach pupal stage (Fig 2A). For VSR, some homozygous individuals reached adulthood (6 males and 1 female) but failed to produce progeny when presented with fertile mates (suggesting that they may be sterile; See Methods). Only ANR, VSK and the triple reversion VNR (i.e. representing the ancestral haplotype) showed viabilities similar to the current ASK haplotype of *D. melanogaster*. Additionally, while the intermediate haplotypes ANR and VSK did not show viability defects, they do exhibit defects in adult locomotion (Fig 2B). Thus, it appears that all three amino acid substitutions are involved in epistatic interactions, and it is apparent that all possible paths from ancestral to the current *D. melanogaster* state would involve transiting through a less fit intermediate state (Fig 2C).

Motivated by these findings, we next asked about the dominance of the deleterious effects associated with intermediate states. To examine dominance effects, we created heterozygotes (x/VNR) carrying one intermediate state (x) and one ancestral haplotype (VNR). We found that all intermediate states have viabilities that are not distinguishable from that of the homozygous VNR or ASK haplotypes (Fig 2A). Similarly, all the intermediate haplotypes lacked locomotion defects when heterozygous (Fig 2B). Together, these results suggest that all deleterious effects associated with intermediate states are recessive. The finding that the deleterious effects of substitutions are recessive has important implications for how we expect this cluster of substitutions to have become established in the *D. melanogaster* lineage. Notably, all paths involving sequential substitution of amino acid substitutions would be deleterious, however, each of these substitutions may have persisted as polymorphisms in the population for some time, assembled into a favorable haplotype and fixed concomitantly (Fig 3). Despite the locomotor defects associated with ANR and VSK (Fig 2B), the fitness effects of these defects may be small. If so, the formation of heterozygotes involving these haplotypes (i.e. x/ANR or x/VSK, where x is any other derived haplotype) could provide additional permissible evolutionary paths. Given this possibility, for a subset of cases we tested whether flies that are heterozygous for two derived haplotypes are viable (Table S2). We find that ASR/ANR and ANK/ANR appear to be perfectly viable, as are VSK/VNK heterozygotes. While we did not test them for fertility, the viability of these genotypes has important implications for how the fitness landscape from VNR to ASK may have been traversed.

**Figure 3.**
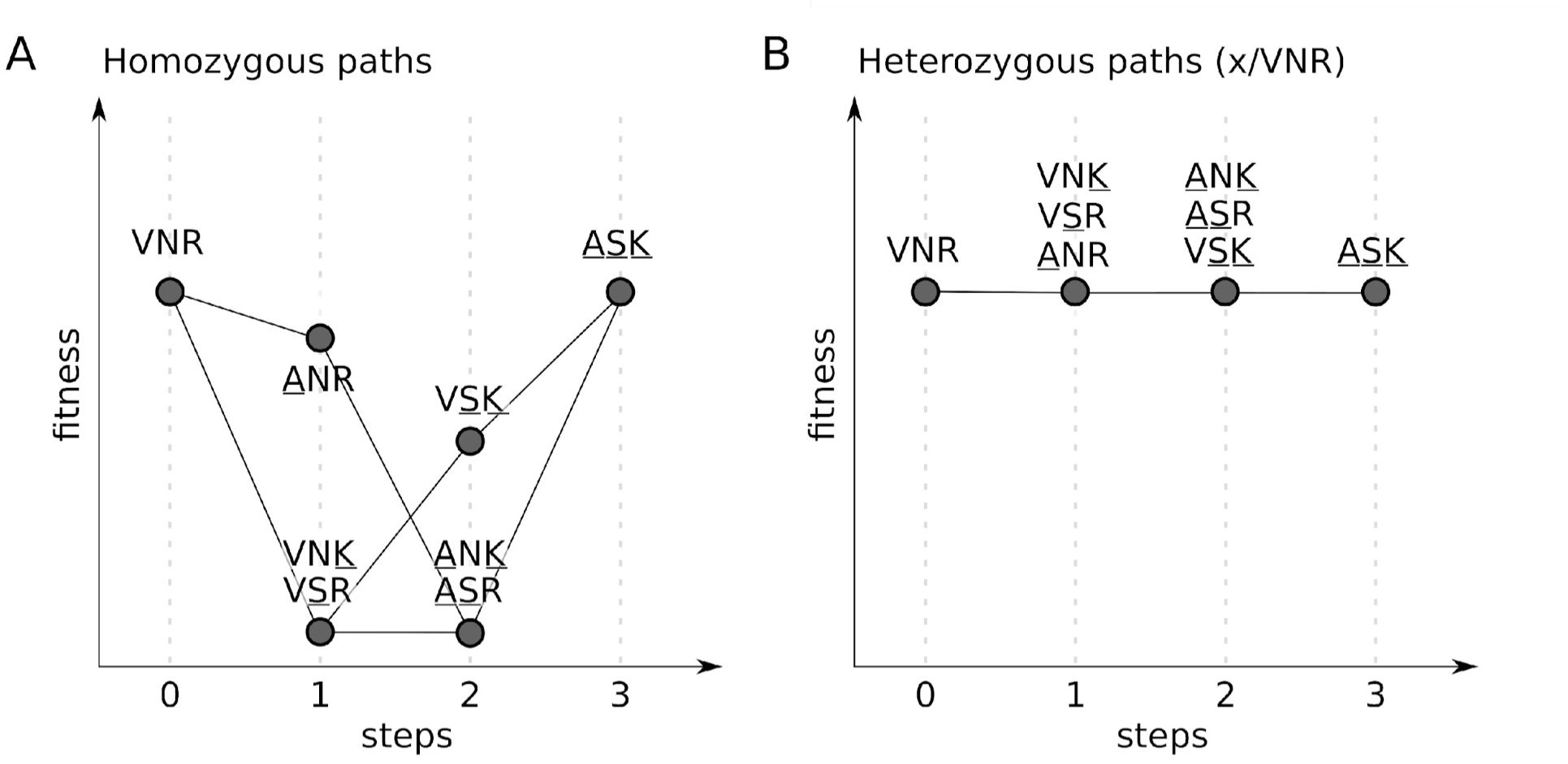
Adaptive paths are only accessible via heterozygous states. Shown is a schematic representation of the fitness landscape for the amino acid substitutions in Trio. The x-axis represents the number of mutational steps from the ancestral haplotype (VNR) to the extant *D. melanogaster* haplotype (ASK). The y-axis represents fitness based on viability and locomotion phenotypes of each haplotype in (A) homozygous *D. melanogaster* flies or (B) heterozygous flies carrying one ancestral VNR haplotype.

As this cluster of substitutions is inferred to be positively selected (Fig 1A), we might expect the ASK haplotype to confer a fitness advantage relative to the ancestral haplotype VNR in *D. melanogaster*. However, the viability and locomotion phenotypes of homozygous VNR flies are not distinguishable from ASK flies. We also find that levels of fertility of the VNR and ASK strains are not distinguishable (Fig S6). This implies that either the ASK haplotype is not strongly beneficial (potentially even neutral) relative to the VNR haplotype, or that a fitness benefit occurs for a phenotype (or in an environment or an ancestral genomic background) that we have not investigated. While we failed to detect a difference in viability and fertility between the VNR and ASK haplotypes, it should be noted that we are underpowered to detect effects that are smaller than 30% given our sample sizes and 10% if our sample sizes were 10-fold larger (Fig S6). In *D. melanogaster* populations, a 1% effect would already correspond to a very strong fitness difference in population genetic terms.

## Discussion

Adaptive protein evolution is widespread in plant and animal genomes and has a major impact on levels of genome-wide variability. Most inferences about this process rest on the assumptions that amino acid substitutions act independently and that the intensity of selection on them remains constant. Increasing evidence, however, suggests that interactions among substitutions—especially those in close physical proximity—may be common. These interactions raise an unresolved question: are clusters of closely linked amino acid substitutions, whether adaptive or not, typically fixed sequentially or simultaneously within a species? Despite its importance, direct empirical evidence for such epistatic interactions among spatially-clustered substitutions remains scarce, particularly in multicellular eukaryotes, where most reports have been anecdotal. To address this gap, we systematically searched for proteins with experimentally tractable clusters of amino acid substitutions that are inferred to be fixed by positive selection. Among the candidates, we identified Trio, a Rho guanine nucleotide exchange factor, as a promising test case. Our analyses suggest that the cluster of three lineage-specific amino acid substitutions in Trio are unlikely to have fixed sequentially along the *Drosophila melanogaster* lineage. Instead, negative epistatic interactions among residues, together with the recessive deleterious effects of the derived variants, depending on epistatic interactions among them, make it more likely that all three substitutions fixed together as a single haplotype. This work provides one of the few direct demonstrations that spatially-clustered substitutions in a multicellular eukaryote can interact epistatically and have measurable effects on whole-organism phenotypes relevant to fitness.

### The utility of *in vivo* studies in Drosophila

Most experimental studies of epistasis have been performed *in vitro*. These approaches are powerful because they allow systematic exploration of epistatic interactions within a protein. For example, deep mutational scanning can efficiently map interactions among sites and reconstruct a protein’s fitness landscape (44,55,56). Such studies have been invaluable for uncovering principles of protein structure–function relationships and for providing quantitative descriptions of higher-order epistasis. However, the phenotypes measured *in vitro*—such as enzymatic activity or ligand binding—are tied to the specific function of the protein being studied and can be difficult to interpret in the context of their effects on organismal fitness.

Moreover, many proteins have functions that are not amenable to straightforward biochemical assays. In contrast, *in vivo* studies – most of which have been carried out in microbial systems (46–48) – provide the opportunity to measure phenotypes more directly relatable to fitness (for example, growth rates). They also provide a way to investigate amino acid substitutions in proteins that are otherwise difficult to assay biochemically, thereby broadening the range of proteins that can be experimentally explored. While microbial systems remain important, key differences between microbes and multicellular diploid organisms have important implications for the mechanistic basis for constraints on adaptation. In particular, multicellular organisms display more complex, higher-order phenotypes—such as those related to development, survival, behavior and reproduction—that in turn increase the complexity of the potential epistatic, pleiotropic and dominance dependencies of adaptive protein evolution. Our study represents one of the few to test the *in vivo* effects of specific adaptive amino acid substitutions on phenotypic traits like viability, fertility, and locomotion in the context of a multicellular organism (see also (75)). Such studies provide a framework for more systematically exploring constraints on adaptive protein evolution in the context of diploid multicellular organisms.

We note, however, that *in vivo* phenotypic analyses are inherently context dependent. While no additional substitutions occur within the DH-PH domain or in the interacting GTPases Rac1 and Rac2 along the *D. melanogaster* lineage, three other amino acid changes did occur elsewhere in the Trio protein. Additionally, we conducted our experiments in a single, contemporary *D. melanogaster* genomic background under laboratory conditions. The deleterious and epistatic effects observed for the intermediate haplotypes are therefore conditional on this specific genetic, genomic and environmental context. It is possible that these intermediates were tolerated in an ancestral genomic background or under alternative ecological conditions. Such caveats are an inherent limitation of all *in vivo* studies.

### Implications for the dynamics of adaptation

Although intramolecular epistasis is recognized as an important factor of protein evolution (52,76,77), how groups of interacting amino acid substitutions arise and reach fixation in functional regions of proteins without compromising fitness remains poorly understood. In principle, epistasis can allow sets of mutations that are individually deleterious or neutral to confer a benefit when combined, yet the evolutionary paths by which such groups of substitutions become established are often opaque. A number of previous studies that have directly investigated the timing and dynamics of epistatic mutation fixation generally support a sequential fixation model (43,45,61,78,79). In such models, mutations fix in a stepwise manner, with each substitution either being neutral or slightly deleterious until the beneficial combination is assembled. This question has largely been unexplored in the context of diploid organisms where dominance relationships of alleles may also play an important role.

For the cluster of amino acid substitutions in Trio, we show that individual substitutions, as well as pairs, are deleterious when present in the homozygous state. This implies that any evolutionary trajectory involving the stepwise, sequential substitution of these amino acids would need to traverse through strongly deleterious intermediate states (Fig 3), making such paths highly unlikely. These results therefore provide strong evidence that this cluster did not arise through a process of sequential fixation. Instead, since these deleterious effects are recessive, we propose that the cluster was more likely to have assembled at the polymorphic phase, with all three substitutions (ASK) eventually fixing concomitantly. Furthermore, ANR and VSK homozygotes are viable (Fig 2), as are ASR/ANR, ANK/ANR and VSK/VNK heterozygotes (Table S3). If other heterozygotes of two derived haplotypes are also viable, this could potentially open additional evolutionary pathways. While waiting times for sequential mutation are expected to be long, the viability of heterozygotes implies that recombination could generate new haplotypes, reducing the time by orders of magnitude (80).

Theoretical modelling suggests that in large populations, deleterious intermediate genotypes can persist at low frequency long enough for a compensatory mutation to arise, restoring fitness and allowing the full combination to spread and reach fixation (81–83). In diploids, recessive lethal haplotypes can persist in the population for many generations (Fig S7), providing an opportunity for a second mutation or a recombination event to occur. This dynamic can lead to the fixation of the final haplotype at a substantially higher rate than the neutral expectation, even when the fitness advantage of the final haplotype is small (81).

Furthermore, if recessive deleterious haplotypes also confer a dominant fitness advantage (a.k.a. a “dominance reversal”), they can be maintained indefinitely and at a moderate frequency (Fig S7), further enhancing the formation of favorable epistatic combinations. This scenario is comparable to a dominance shift following an environmental change described by (84). Consistent with this view, several recent examples of adaptive protein evolution have documented dominance reversals, where adaptive mutations exhibit dominant beneficial effects but recessive pleiotropic costs. Such a pattern has been observed in the evolution of Na+,K+-ATPase resistance to cardiotonic steroids (58), GABA receptor resistance to terpenoids (85), and acetylcholinesterase resistance to organophosphates (86). The dominance of the beneficial effects associated with adaptive substitutions at Trio is not known, but given the recessivity of deleterious intermediate states, it may represent a similar example of adaptive protein evolution.

Notably, the scenarios described above assume that mutations arise independently. An alternative way to traverse fitness valleys caused by deleterious intermediates is that all three substitutions (or two of the three) arose simultaneously in a single mutational event. Multi-nucleotide mutational events have been documented in several systems (87–91), and such events could facilitate the coordinated emergence of interacting amino acid changes. Based on Drosophila mutation-accumulation lines and human parent-offspring trios, multi-nucleotide mutational events account for about 3% of the *de novo* mutations in the germline across eukaryotes (88,91). While this mechanism could, in principle, account for the pattern observed in Trio, it seems unlikely that a single mutational event would generate multiple substitutions that are individually deleterious but jointly neutral or beneficial. In any case, multi-nucleotide mutational events may contribute to the clustered patterns observed in some of our other candidate genes. Notably, the documented excess of spatially clustered amino acid substitutions in Drosophila proteins persists between substitutions separated by introns, suggesting that multi-nucleotide mutation alone cannot fully explain the observed clustering pattern (34). Thus experimentally examining clusters of substitutions that are separated by introns may help distinguish substitutions that arose independently from those generated by a single mutational event.

Further, although our candidate clustered substitutions are inferred to have been fixed by positive selection using a McDonald-Kreitman-base approach, this signal does not distinguish between strict adaptation—where the derived haplotype confers a fitness advantage relative to the ancestral state—and compensatory evolution. Several studies have highlighted genome-wide signatures of compensatory protein evolution (33,36,77,92,93). Compensatory evolution can also produce clustering patterns, as compensatory substitutions are often located closer to the focal mutations (33,36).

While some of our candidates for proteins with putatively adaptive clusters of amino acid substitutions may be due to compensatory interactions, compensatory dynamics and positive selection are not mutually exclusive hypotheses. In the case of Trio, given the severely deleterious states of most intermediates, it seems unlikely that the fitness of the extant haplotype ASK is effectively neutral with respect to fitness. Despite this, based on the current data, neutral fixation cannot be ruled out. Importantly, even if many of the clusters we identified, including in Trio, are compensatory rather than adaptive, they remain highly informative. Understanding how such interacting substitutions fix—and how they shape patterns of molecular variation—is critical for interpreting population genetic signals of selection and for clarifying the mechanisms underlying protein evolution.

### Implications for the interpretation of population genetic inferences

Over the past several decades, population genetics has undergone a paradigm shift in how we understand the role of positive selection in protein evolution and its impact on genomic variation in plants and animals (8,12). In Drosophila, estimates suggest that 50–90% of amino acid divergence exceeds neutral–deleterious model predictions and is therefore attributed to positive selection (20,25,94,95). Independent support for adaptive substitutions comes from hitchhiking signatures near individual amino acid substitutions (18), but these analyses tend to yield somewhat lower estimates of the proportion of divergence that is adaptive.

If, however, epistasis among amino acid substitutions (as observed for Trio) is common in protein evolution, the McDonald–Kreitman framework may overestimate the proportion of adaptive divergence (96). This is because the framework assumes independence among sites and a constant direction and strength of selection over time, both of which are violated when substitutions interact epistatically. Although this issue has been raised conceptually, the consequences of pervasive epistasis for interpreting McDonald–Kreitman results remain largely unexplored (96).

More broadly, widespread non-independence among amino acid substitutions could alter how we interpret their effects on genome-wide diversity. In standard models of genetic hitchhiking, reductions in linked neutral diversity are attributed to substitutions fixing independently (18,97,98). However, intuitively, if several closely linked substitutions fix simultaneously rather than sequentially, the apparent average strength of selection per substitution inferred from local reductions in linked diversity would be underestimated, since its impact would be distributed across multiple fixed sites. How such non-independence—particularly in light of the dominance and epistasis uncovered in the case of Trio—affects inferences of population genetic parameters is an important open question for future research.

### Conclusions

By studying a cluster of putatively adaptive amino acid substitutions in the *D. melanogaster* protein Trio, we uncovered surprisingly strong deleterious epistatic interactions among residues of this cluster. Because these deleterious effects are recessive, they are more likely to have persisted in the population as polymorphisms before combining into a beneficial haplotype that eventually became fixed in the species. This may be a common phenomenon in adaptive protein evolution and has important implications for how we interpret its associated population genetic signatures. This case study lays the groundwork for more systematic *in vivo* studies of putatively adaptive amino acid substitution, where the fitness effects of mutations can be measured at multiple levels, including whole-organism phenotypes related to fitness. Such studies will help clarify how epistasis, pleiotropy, and dominance interact to shape protein evolution, deepening our understanding of the evolutionary constraints that govern evolutionary trajectories in multicellular diploid organisms.

## Methods

### Identification of adaptive clusters

We generated multiple sequence alignments of orthologous proteins using *D. melanogaster* ISO1 (GCA_000001215.4), *D. simulans* w501 (GCA_016746395.2), *D. yakuba* NY73PB (GCA_016746365.2), *D. santomea* STO CAGO 1482 (GCA_016746245.2) and *D. teissieri* GT53w (GCA_016746235.2) One-to-one orthologs were identified using a reciprocal best-hit exonerate approach (35,99). Predicted *D. melanogaster* (strain ISO1) proteins were aligned to each genome sequence to extract candidate proteins, and each extracted protein was subsequently realigned to its corresponding *D. melanogaster* protein. The longest predicted transcript for each protein in *D. melanogaster* was used to represent each protein and multispecies alignment was created using PRANK (-codon parameter, v.170427; (100)). Ancestral sequences were inferred based on this alignment and the species tree using PAML’s baseml function (v 4.9; (71)) with the following parameters: model=7, kappa=1.6, RateAncestor=2. MACSE (- max_refine_iter 1 parameter, v. 2.07; (101)) was then used to align sequences from the *D. melanogaster* Zambia population (n = 82; (70)) with the reconstructed *D. melanogaster*–*D. simulans* ancestral sequence.

To infer selection on protein sequences, we applied Model Averaged Site Selection via Poisson Random Field (MASS-PRF), which relies on both polymorphism and divergence data (26). In this framework, sequences from the Zambia population served as polymorphism data, while the reconstructed *D. melanogaster*–*D. simulans* ancestral sequence was used as the outgroup to quantify lineage-specific divergence. To prepare the input files required for running MASS-PRF (26), we generated consensus sequences capturing polymorphism and divergence changes for each protein, as follows. To create the polymorphism consensus sequence, monomorphic codons were replaced by “*”, synonymous codons by “S” and non-synonymous changes were replaced by “R”. To create the divergence consensus sequence, outgroup codons that were present in the ingroup sequences were replaced by “*”, outgroup codons were replaced by “S” or “R” if the ingroup codon was fixed and the outgroup codon was synonymous or non-synonymous to it respectively. If the ingroup codon was polymorphic and the outgroup codon was different from both of these, the codon was replaced by “-”. For proteins longer than 900 amino acids, Consensus sequences were split into fragments of 900 codons or smaller.

MASS-PRF was run on each protein fragment using parameters –ic 1 –sn 82 -o 1 -r 1 - ci_r 1 -ci_m 1 -s 1 -exact 0 -mn 30000 -t 2. Of 9137 proteins analyzed, MASS-PRF ran successfully for 5355 for at least one fragment per protein (Across all fragments, 5743: successful run; 4082: too little substitution information to run; 748: out of memory). Successful runs were screened for positive selection. A site was estimated to be under positive selection if the lower 95% confidence bound for gamma > 4, which corresponds to a false positive rate <0.1 (26). Of the 5355 proteins for which runs were successful, 1144 had at least one site under positive selection. Among them, 919 harbors clustered substitutions, defined by at least two sites less than 20 codons apart. Within this set, we found that 679 proteins contain at least one potential experimentally tractable cluster, corresponding to a total of 795 clusters. We defined a cluster as experimentally tractable if exactly 2 or 3 sites that show divergence and positive selection are less than 20 codons apart from each other.

Predicted functional domains (Fig 1 and Fig S1) were found in FlyBase.org and AlphaFold predictions are taken from the AlphaFold Protein Structure Database (EMBL-EBI). Superimposition of the AlphaFold prediction and the crystal structure (when available) was performed using Mol* Viewer (102). For protein stability analyses (Table S1), structures of the *D. melanogaster* Trio DH-PH domain, as well as the ancestral and all the intermediate versions of the domain were predicted using AlphaFold2 and MMseqs2 implemented in ColabFold v1.5.5 (103) using default parameters. The effects of each single mutation on stability (ΔΔG) were predicted using Dynamut2 (72), ThermoNet based on Rosetta modules (73,104,105) and ACDC-NN (istruct function) (74).

### CRISPR/cas9-mediated editing: Plasmid assembly

Genomic DNA was isolated from ywISO8 flies using a “squish prep” protocol (106). Homology arms were amplified with PCR from this genomic DNA using Q5 polymerase from NEB (Table S2). The plasmid backbone and dsRed were amplified from plasmid: pScarlessHD-DsRed, a gift from Kate O’Connor-Giles (Addgene #64703) (Table S2). The four fragments were combined at an equimolar concentration (150 fmoles each) and assembled using NEBuilder HiFi DNA assembly kit. To verify that assembly was successful, PCR was performed using primers that spanned multiple fragments. The NEBuilder HiFi DNA assembly kit result was used to transform NEB 5-alpha competent E. coli. To verify that the transformation was successful, colony PCR was performed on individual bacteria colonies using pairs of primers that spanned multiple fragments using LongAMP or regular Taq. Final plasmid assembly was verified by a Tn5 tagmentation-based Illumina sequencing protocol (107) followed by *de novo* assembly. Plasmids were extracted using Qiagen’s QIAprep Spin Miniprep kit.

### CRISPR/cas9-mediated editing: Site-Directed Mutagenesis

Starting from a base plasmid sequence representing the current *D. melanogaster* haplotype (ASK), we used site-directed mutagenesis (SDM) to create all possible intermediates leading back to the ancestral A1315V/S1318N/K1335R (VNR) haplotype. To do this, we used the Agilent QuikChange Lightning Multi Site-Directed Mutagenesis Kit in two reactions: one containing three primers: 1) A1315V, 2) S1318N, and 3) K1335R, and one containing two primers: 1) A1315V, S1318N and 2) K1335R. The standard Agilent kit protocol was followed. First, PCR was performed without the addition of QuikSolution, as it was not necessary. This was followed by DpnI digestion (provided) and transformation into the provided competent cells. Colony PCR using Taq polymerase was performed on 48 bacterial colonies from each reaction (Table S2). Amplicons from each sample were barcoded with a unique combination of Illumina-compatible i5 and i7 adapters and sequenced to assess the presence of the desired mutations. Plasmids were extracted using the Qiagen Plasmid Midi Kit for CRISPR injection. The design for mutation 3 (K1335R) is completely scarless. All the other combinations introduce a single synonymous single nucleotide polymorphism that is naturally present within *D. melanogaster* populations. Plasmids were obtained for the following combinations: A1315V (VSK), A1315V/S1318N/K1335R (VNR), A1315V/S1318N (VNK), S1318N (ANK), K1335R (ASR), S1318N/K1335R (ANR). One additional haplotype, A1315V/K1335R (VSR) was generated using a second SDM reaction starting from A1315V plasmid to introduce K1335R.

### CRISPR/cas9-mediated editing: injections and post-injection processing

gRNAs were created from the shared tracrRNA and a site-specific crRNA, ordered as two separate Alt-R® CRISPR-Cas9 RNA fragments from IDT (Table S2). For each gRNA, tracrRNA and crRNA were annealed by adding 2 μl of crRNA at 100 μM, 2 μl of tracrRNA at 100μM, and 2 μl of IDT nuclease-free duplex buffer and incubating at 95 °C for 5 minutes, then allowing the mixture to come to room temperature on the benchtop. gRNA was loaded into the Alt-R® S.p. Cas9 Nuclease from IDT to make the ribonucleoprotein (RNP) complex: First, stock Cas9 solution was diluted into an equivalent volume of IDT nuclease-free duplex solution. Then 2.68 mL of both annealed gRNAs and 1.8 mL of the diluted Cas9 protein were mixed and incubated at room temperature for 5 minutes. Finally, 7.5 mg of donor plasmid was added, and the volume was increased to 30 mL with water. CRISPR-cas9 injections were done in line “ywISO8” to facilitate ease of fluorescent screening. Injected flies were crossed to a double balancer line (BL#2537: w[*]; TM3, Sb[1] Ser[1]/TM6B, Tb[1]). Progeny of the crosses were screened for red fluorescent eyes and mated with siblings with the same balancer chromosomes to establish a stable line. pBac excision was performed following (108): DsRed embryos were injected with 250ng/mL phsp-pBac plasmid (109), heat shocked 1 hour after injection for 1 hour at 37 C. Injected flies were then crossed with a balancer line (Bloomington #23232: w[*]; ry[506] Dr[1]/TM6B, P{Dfd-GMR-nvYFP}4, Sb[1] Tb[1] ca[1]). Progeny of the crosses were screened for non-fluorescent eyes and sib mated to establish mutant lines.

### CRISPR/cas9-mediated editing: Background homogenization

In order to fully control for genomic background effects, two balancer lines with a fully homozygous genome were created using the inbred line RAL386 (BL#28192; (78) and the chromosome balancer of respectively BL#23232 (w[*]; ry[506] Dr[1]/TM6B, P{Dfd-GMR-nvYFP}4, Sb[1] Tb[1] ca[1]) and BL#35524 (w[1118]; DCTN1-p150[1]/TM3, P{w[+mC]=sChFP}3, Sb[1]). RAL386 males were crossed with virgin females of either balancer line BL#23232 or BL#35524. An F1 male was then backcrossed with a RAL386 virgin female. Resulting fluorescent male progeny were then backcrossed to RAL386 virgin females. Several days after the crosses, the father of each cross was collected and genotyped for chromosome 2 by PCR and Sanger sequencing of a region with a known diagnostic single nucleotide polymorphism. Crosses for which the father was homozygous for RAL386 haplotype for chromosome 2 were kept (Table S2). Fluorescent virgin female progeny from this cross were then backcrossed to a RAL386 male. The balancer chromosomes in the established lines were maintained by screening for fluorescence every 2-3 weeks. *Trio* mutant lines and yw-ISO8 line, used as the control ASK, were crossed alternatively with the two RAL386 balancer lines following the same crosses scheme as described below so the final lines could be balanced with TM6B, P{Dfd-GMR-nvYFP}4, Sb[1] Tb[1] ca[1].

To confirm the genomic background of engineered lines, 10 female individuals were collected from each line and frozen at -20°C. Whole DNA was extracted from each fly individually using Quick-DNA 96 Kit (Zymo research). DNA from each fly from the same line was then pooled. Tn5-tagmentation libraries (107) were prepared and indexed with one specific index for each line and enriched by 12 PCR cycles (HS One Taq, NEB). Libraries were then pooled and size selected for fragments around 500nt using AMPure XP (Beckman Coulter) following manual. The pooled libraries were then sequenced with paired-end 150nt on NovaSeq X Plus 10B Illumina 2x150 (Admera Health). Illumina adapters were trimmed using trim-galore (-q 0, -e 0.1) (v 0.6.7) and reads were aligned to the *D. melanogaster* reference genome (GCA_000001215.4) using bwa mem (v 0.7.17-r1188, (110)). Duplicates were marked with picard (v 2.18.29, Broad Institute) and reads were re-aligned with gatk3 (v 3.8-1-0, Broad Institute). Variants were called and pseudo references generated using samtools (v 1.6) and bcftools (v 1.9) and the PseudoreferencePipeline scripts of the YourePrettyGood pipeline (https://github.com/YourePrettyGood/). Average pairwise divergence (Dxy) between engineered strains and RAL386, yw iso8 and RAL59 was calculated for 1 Mbp non-overlapping windows using RandomScripts#calculatepolymorphismcpp of the YourePrettyGood pipeline (https://github.com/YourePrettyGood/) (Fig S5).

### Drosophila rearing

All fly strains were reared in bottles or vials at 21°C, 50% in day-night cycle of 12hr-12hr on fly food containing agar (0.56%), cornmeal (6.71%), inactivated yeast (1.59%), soy flour (0.92%), corn syrup (7.0%), propionic acid (0.44%), and Tegosept (0.15%) (LabExpress).

### Viability assays

Viability of homozygous strains was measured as the relative survival of homozygous YFP-progeny relative to YFP+ progeny produced by parents balanced with TM6B, P{Dfd-GMR-nvYFP}4, Sb[1] Tb[1] ca[1]. For 1^st^ instar larva, virgin males and females [YFP+, Sb+, Tb+] were crossed and allowed to lay eggs on grape-juice agar plates with yeast paste for 24hr. After 48hr, the yeast paste was washed out through a cell strainer to collect and count fluorescent and non-fluorescent 1st instar larvae. For pupal and adult stage, females were allowed to lay eggs on regular fly food vials for 7 days. [Tb+] and [Tb-] pupae were counted after about 10 days. [Sb+] and [Sb-] adults were counted after about 20 days. We measured the viability of heterozygous individuals the same way except that homozygous VNR virgin females were crossed to males balanced with TM6B, P{Dfd-GMR-nvYFP}4, Sb[1] Tb[1] ca[1]. 60 to 300 larvae, 100 to 200 pupae and 100 to 150 adults were counted for each engineered strain. For VSR, we obtained 6 adult homozygous males and 1 adult homozygous female. To estimate the fertility of those homozygous adult individuals, males were crossed individually with one homozygous VNR virgin female and the female was crossed with one heterozygous VSR male. No progeny were observed after 21 days.

### Locomotion performance assay

We quantified locomotor performance by quantifying startle-induced negative geotaxis, an innate behavior where flies tend to climb against gravity after being mechanically agitated (111). Ten 1-day-old males were grouped with ten females into vials for 7-10 days. Before the assay, the males were transferred into ethanol-cleaned assay tubes, consisting of two tubes attached with tapes with a line set at 4.5 cm from the bottom of the tube. Flies were let to recover for 40 to 60 min before the start of the experiment. The assay tube was tapped 5 times on a fly pad so that all flies fell to the bottom of the tube, and the flies that reached the line in the following 10 sec were counted. Each group of flies were assayed three times, with 5 min recovery between each time, the average of the three replicates was used. The same assay tubes were then assayed the same way after 10 min recovery but counting flies that reached the line in the following 5 sec instead of 10. Four to 14 replicates were performed for each haplotype.

## Acknowledgements

Thanks to N. Okami for help with fly stocks and K. O’Connor-Giles for sharing reagents. Thanks to the Andolfatto, Przeworski and Sella labs for useful discussions. This work was funded by the National Institutes of Health R01 GM115523 to P.A.

## Data availability

Raw data for functional assays and MASS-PRF pipeline and results have been deposited to Github (https://github.com/fborne2/Trio_epistasis/).

