## Supplementary Materials for "Epistasis among clustered lineage-specific amino acid substitutions in the Drosophila Trio protein"

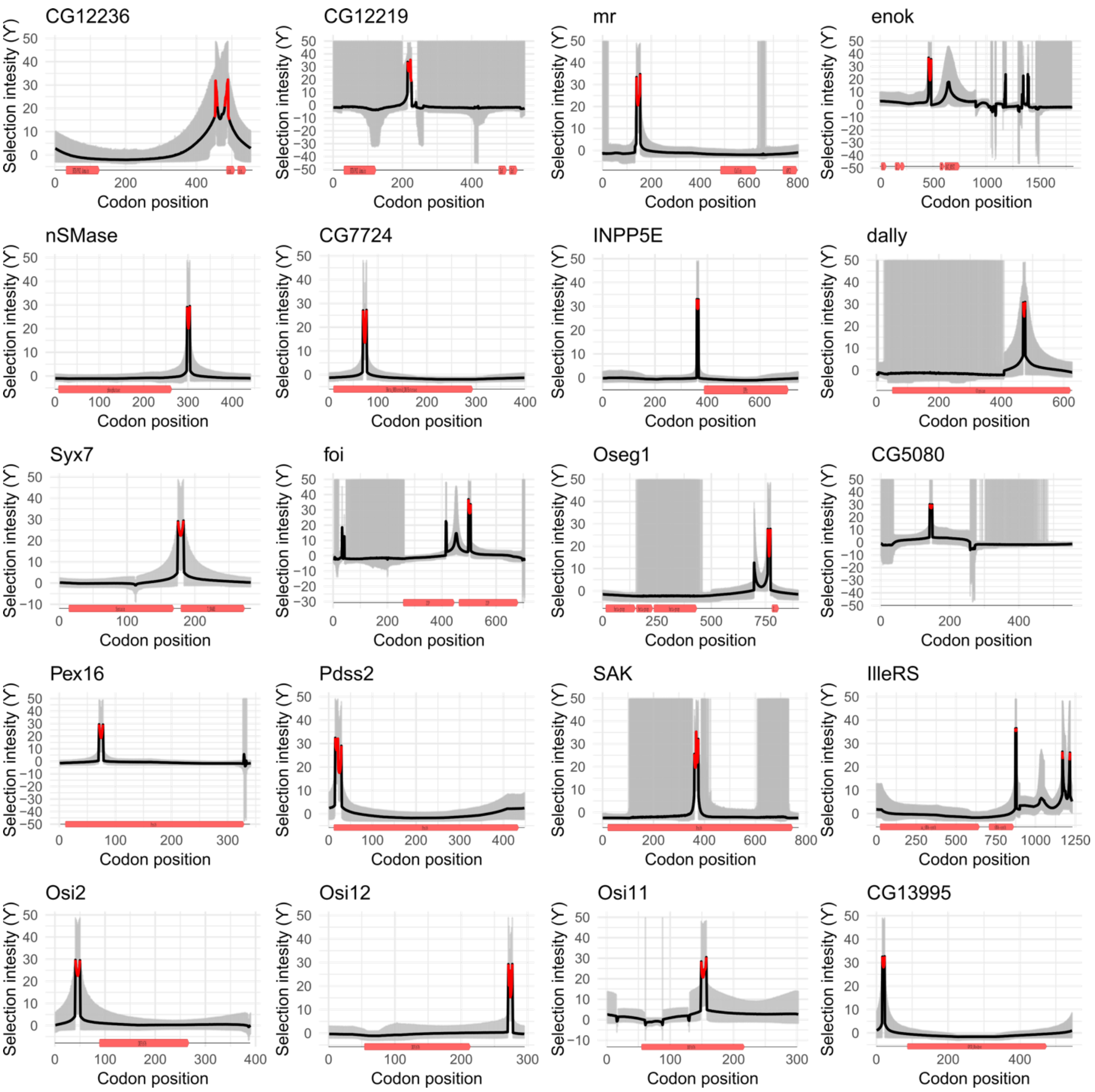


**Figure S1. Additional examples of Drosophila proteins exhibiting lineage-specific clusters of adaptive amino acid substitutions.** Plotted are profiles of selection intensity ($\gamma = 2Ns$) across 20 D. melanogaster proteins inferred with MASS-PRF (1). The black line corresponds to the model-averaged $\gamma$ and the grey areas indicate 95% model uncertainty interval. Red lines indicate regions for which the 95% lower bound of $\gamma > 4$, which corresponds to a false positive rate of <0.1 (see Methods). Protein models are represented below each selection intensity profile. Functional domains are represented by red boxes with their names in black.

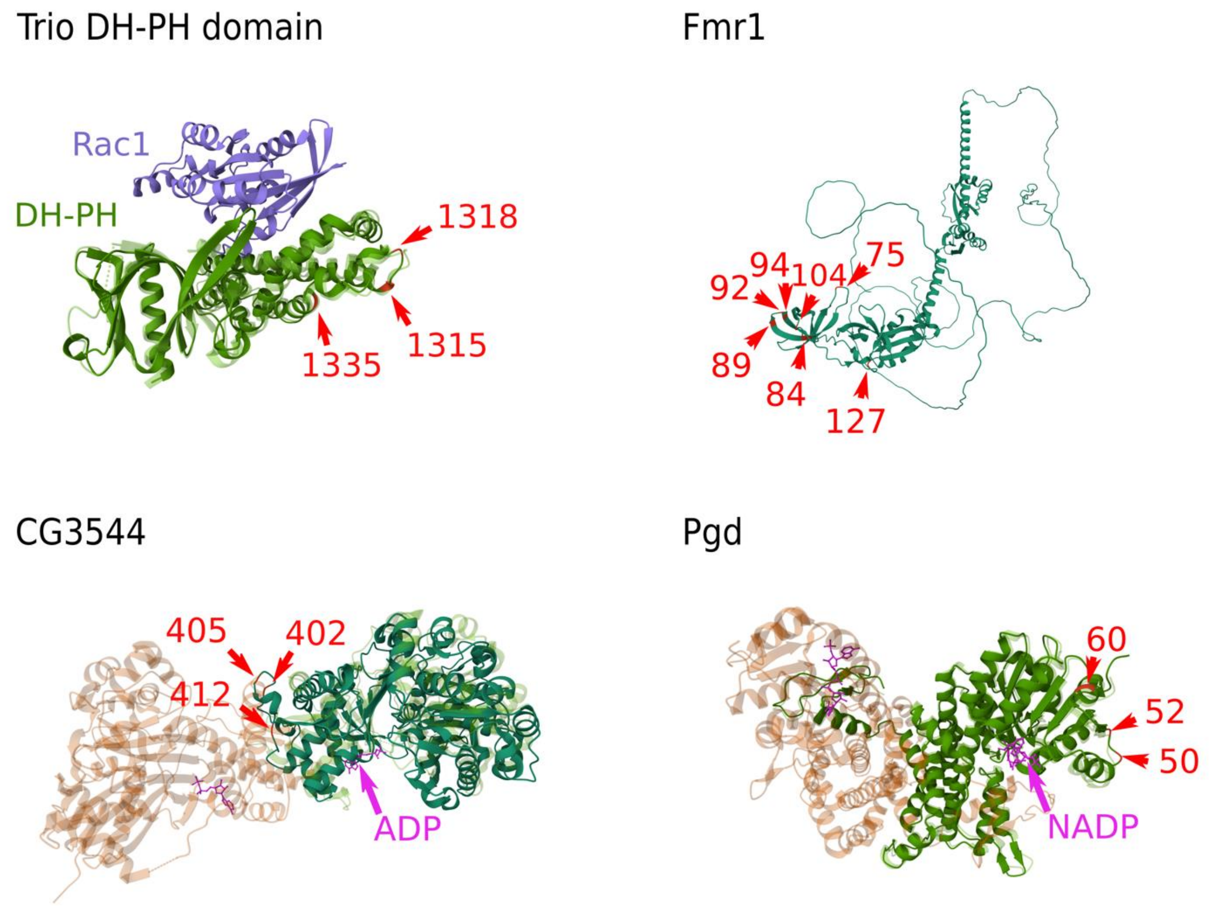


**Figure S2. Examples of Drosophila proteins exhibiting lineage-specific clusters.** AlphaFold predictions of the 3D structure of the *D. melanogaster* proteins superimposed on to the available homologous PDB structures (See Fig 1 for more details).

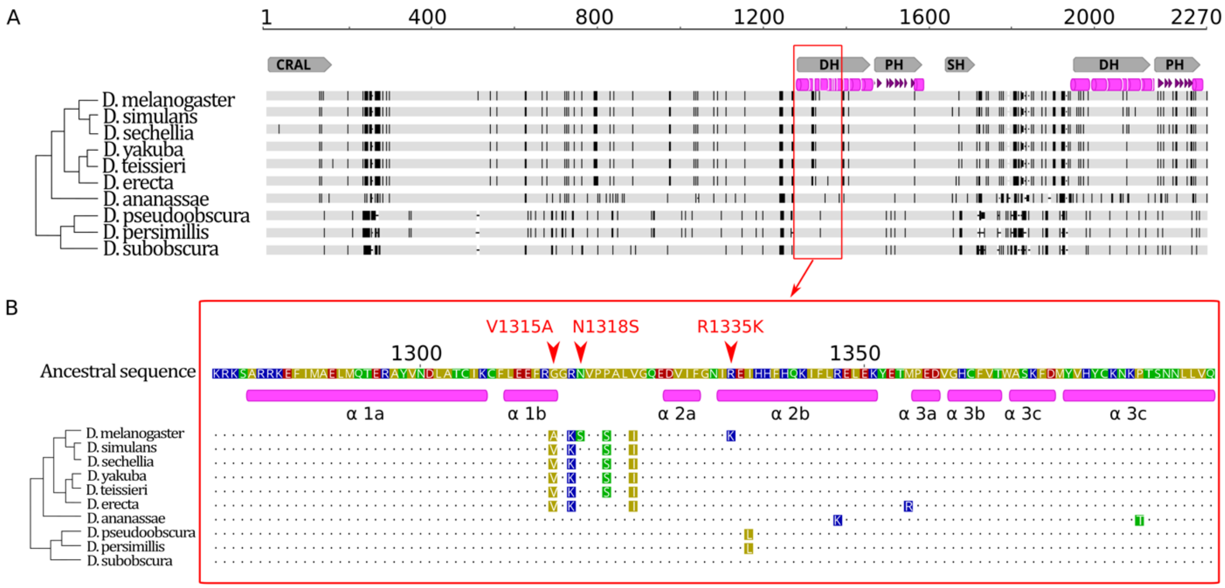


**Figure S3. Trio protein alignment within Drosophila genus.** (A) Alignments of the full protein sequence with large indels removed. Disagreements to ancestral sequence are represented in black and agreements in grey. functional domains are represented by grey boxes. (B) Focus on the Trio cluster region. Amino acids are colored according to their polarity (yellow: non-polar; green: polar; red: polar, acid; blue: polar, basic). Dot represents agreements to the ancestral sequence. Positions of the helix are based on the human structure (PDB 7SJ4, (2)). Species tree from (3,4). Branches are not at scale. Muscle alignment (v 3.8.425) implemented in Geneious Prime (v 2019.0.4). Trio *D. melanogaster-D. subobscura* ancestral sequence has been estimated using codeml function of PAML (v 4.9; (5)) using protein sequences of *D. melanogaster*, *D. simulans*, *D. yakuba*, *D. ananassae*, *D. pseudoobscura*, *D. willistoni*, *D. virilis* and the corresponding species tree.

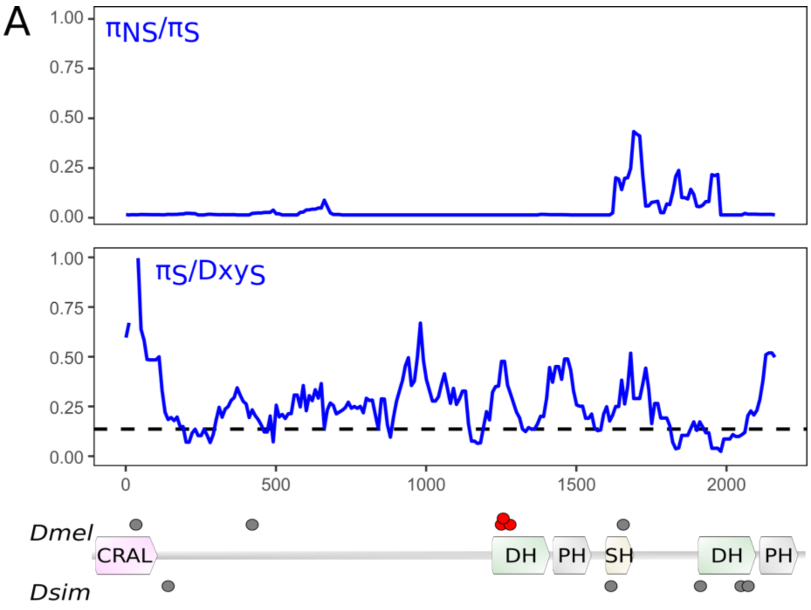


B

| McDonald-Kreitman test | | | | | |
|---|---|---|---|---|---|
| Pop | Pn | Ps | Dn | Ds | Fisher's exact test |
| ZI | 41 | 375 | 13 | 88 | P=0.468 |
| EF | 15 | 144 | 13 | 94 | P=0.543 |
| RAL | 97 | 140 | 13 | 22 | P=0.7157 |
| SD | 16 | 216 | 13 | 95 | P=0.1435 |

C

| null vs branch-site model | | | | |
|---|---|---|---|---|
| Model | $\ell$ | fp | df | $2\Delta\ell$* |
| Dmel branch-site null | -16037.8599 | 22 | 1 | 3.1160 |
| Dmel branch-site | -16036.3019 | 23 | | |

* critical value is $\chi^2 = 3.84$ at 5% significance

**Figure S4. Classic tests of selection do not show sign of positive selection in Trio.** (A) population genetic statistics do not show sign of positive selection ($\pi_{NS}/\pi_S$) or selective sweep ($\pi_S/Dxy_S$). S: synonymous; NS: nonsynonymous. Diversity π and divergence Dxy have been calculated in sliding windows of 100 codons with 10 codon steps using SNPGenie (6). π has been calculated using individuals from the Raleigh population (n=210) (7) and divergence Dxy has been calculated using the *D. melanogaster* reference sequence (GCA_000001215.4) and the *D. simulans* reference sequence (GCA_016746395.2). Black dashed line corresponds to the average value of pi_syn4f / Dxy_4fsyn for the middle of chromosome arm 3R in the Raleigh population (8). (B) McDonal-Kreitman tests performed using DNAsp6 (9) on several *D. melanogaster* populations are not significant. ZI: Zimbabwe; EF: Ethiopia; RAL: Raleigh; SD: South Africa (7). (C) Branch-site model performed with PAML (v 4.9; (5, 10)) using the Drosophila species tree does not show sign of selection in the *D. melanogaster* branch. $\ell$ : log-likelihood score.fp: free parameters; df: degree of freedom; LTR statistics $2\Delta\ell = 2(\ell_1 - \ell_0)$.

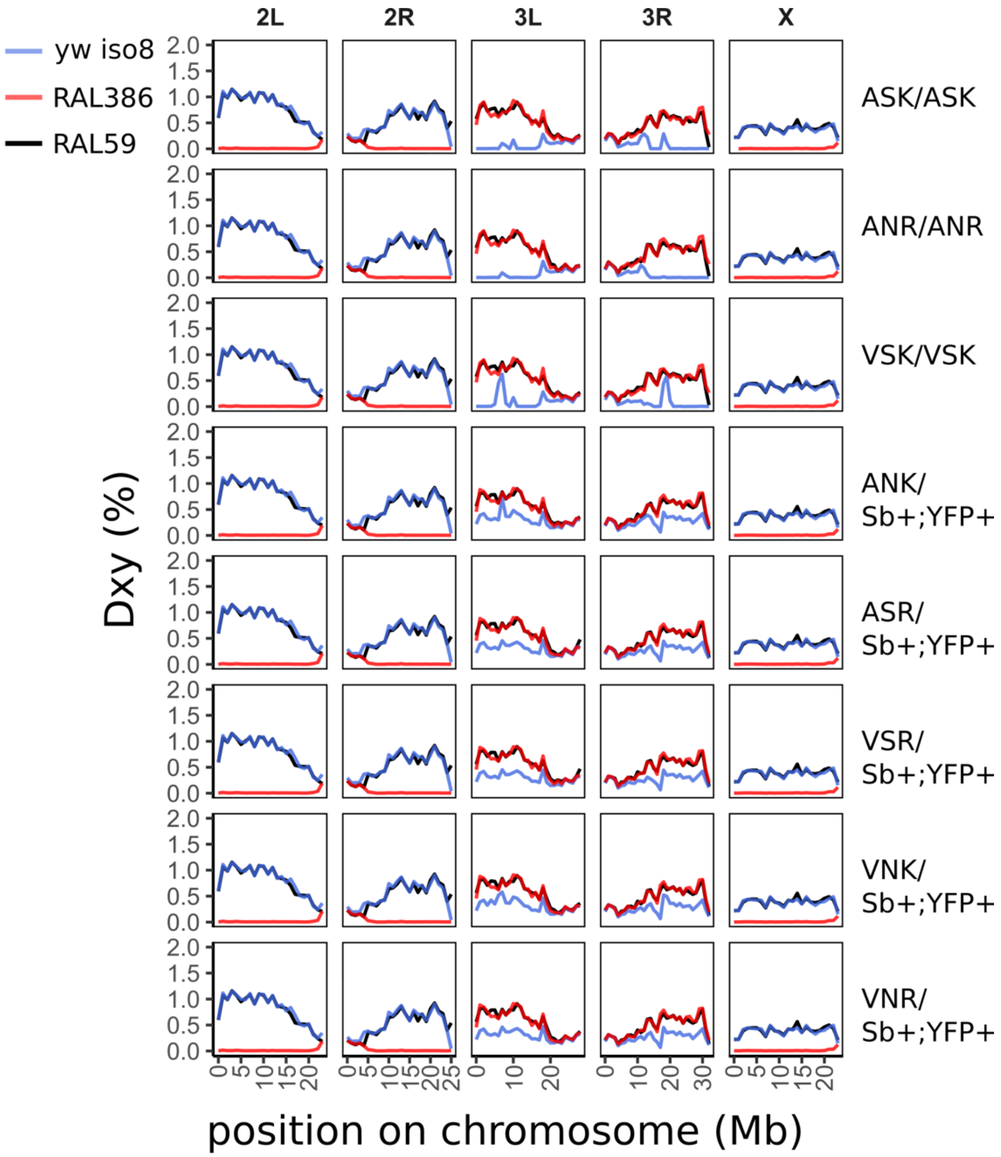


**Figure S5. Sequencing confirmation of engineered lines.** Average pairwise divergence (Dxy) between engineered strains and RAL386 (red line), yw iso8 (blue line) and RAL59 (black line) was calculated on 1 Mbp non overlapping windows along chromosome arms (See Methods).

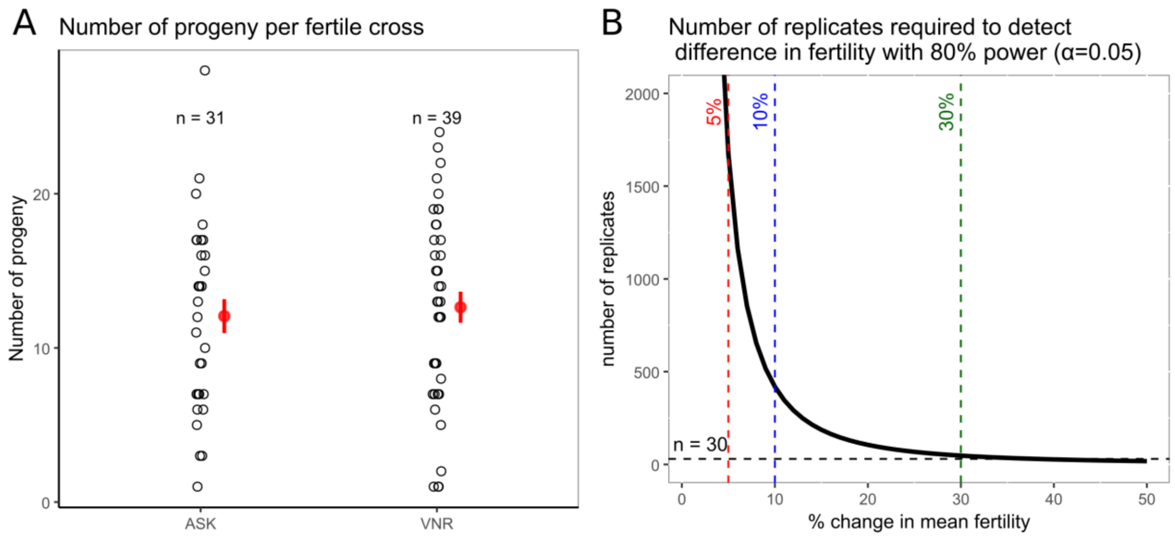


**Figure S6. Fertility of homozygous ASK and VNR strains.** (A) Single 1-3 day-old virgin females were crossed with single 1-3 day-old virgin males from the same line and allowed to mate for 24hr. Adult offspring were counted 20 to 27 days later. Each dot represents the number of progeny for one fertile cross. Red circles and bars represent the mean and standard error, respectively. The two haplotypes do not show a significant difference (t-test, p=0.8961). (B) Curve shows the number of replicates required to detect a given difference in mean fertility between genotypes with 80% power ($\alpha$ =0.05). This calculation is based on the pooled standard deviation of progeny counts estimated from the ASK and VNR strains. The dashed red and blue and green vertical lines indicate 5%, 10% and 30% differences in mean progeny, respectively. The horizontal dashed green line indicates the current experimental sample size (n ~ 30). Although 1-5% change in fertility can contribute greatly to selection, experimental design to detect this change would require >1500 replicates. Script is available at https://github.com/fborne2/Trio_epistasis/.

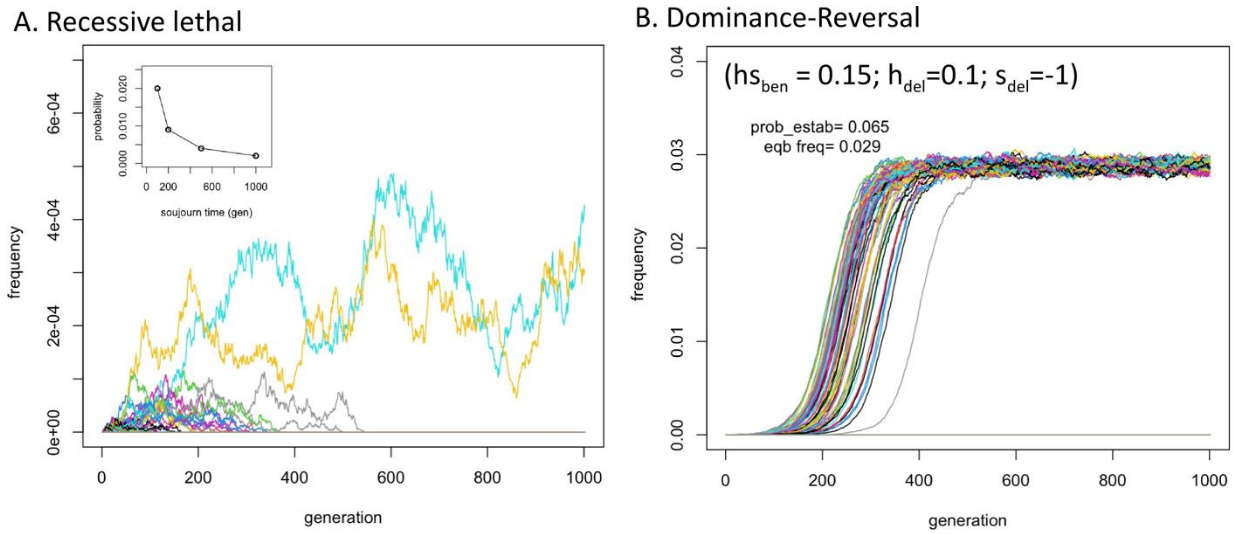


**Figure S7. Example dynamics of recessive lethals and dominance-reversal scenarios.** (A) Forward simulation of newly arising fully recessive (h=0) lethal mutations for a population size of $10^6$. The inset indicates the estimated probability that a mutation will persist in the population for given sojourn time thresholds. Noteworthy is that newly arising lethals can persist in large populations, albeit at very low frequencies, for an appreciable number of generations. Carter and Wagner (2022) show that in large populations like this, for modest mutation rates to secondary mutations that render the haplotype strongly beneficial, the probability of fixation can substantially exceed the neutral expectation (11). (B) Forward simulation (population size of $10^6$) for newly arising mutations that exhibit a "dominance reversal": i.e. in this case, a partially recessive lethal ($h_{del}$=0.1) that has dominant beneficial effects ($hs_{ben}$=0.15) on another fitness-related phenotypic axis. The resulting expected trajectory is the classic expectation for a balanced polymorphism. Prob_estab is the estimated probability of such an allele becoming established in the population, and eqb_freq is the equilibrium frequency conditional on the mutation becoming established. Modest deviations from analytical approximations are likely due to the large selection coefficients being modelled. Noteworthy is that, conditional on becoming established, such mutations persist indefinitely at modest frequencies, and long enough for secondary mutations that render the haplotype neutral or beneficial. In the case of a secondary mutation that renders the haplotype neutral, the probability of fixation will be approximately equal to the equilibrium frequency and substantially higher if net beneficial. Script is available at https://github.com/fborne2/Trio_epistasis/.

**Table S1. Predicted effects of single mutations on protein stability.** The effects of each single mutation on stability (ΔΔG) were predicted using Dynamut2 (12), ThermoNet based on Rosetta modules (13-15) and ACDC-NN (istruct function) (16). ΔΔG is defined as ΔΔG (kcal/mol) = ΔG(mutant) − ΔG(wild-type). All ΔΔG values lie within the interval of –1 to +1 kcal/mol, a range generally considered to be functionally neutral (17,18).

| Change | Dynamut2 | ACDC-NN (istruct) | ThermoNet |
|---|---|---|---|
| **VNR -> ANR** | 0.30 | 0.20 | 0.73 |
| **VNR -> VSR** | -0.27 | -0.0015 | -0.014 |
| **VNR -> VNK** | 1.00 | 0.36 | -0.058 |
| **ANR -> ASR** | -0.28 | -0.019 | -0.11 |
| **ANR -> ANK** | 0.87 | 0.40 | -0.20 |
| **VSR -> ASR** | 0.27 | 0.20 | 0.24 |
| **VSR -> VSK** | -0.86 | 0.40 | -0.18 |
| **VNK -> ANK** | 0.33 | 0.20 | 0.70 |
| **VNK -> VSK** | -0.27 | 0.40 | 0.22 |
| **ASR -> ASK** | 0.85 | 0.36 | -0.13 |
| **ANK -> ASK** | -0.29 | -0.0069 | 0.044 |
| **VSK -> ASK** | 0.47 | 0.20 | 0.75 |

**Table S2. Sequences of primers and other oligonucleotides used in this study.**

| Primers for PCR/Gibson assembly | | |
|---|---|---|
| **Assembly pieces** | **Forward** | **Reverse** |
| Backbone | ttcttgcagtgttAGAAGACCATATACGTCTC | gatgtgtgcgtctGCCCCGAAGACACTATAG |
| LHA | gtgtcttcggggcAGACGCACACATCGGCGG | tttctagggttaaAGCAAGGTATTACTACTATCGATG |
| dsRed | gtaataccttgctTTAACCCTAGAAAGATAATCATATTG | gtataaaaacgtaTTAACCCTAGAAAGATAGTCTGC |
| RHA | tttctagggttaaTACGTTTTTATACCTTACAGATTTGG | tatatggtcttctAACACTGCAAGAAGTTTC |

| PCR conditions for the above primers | | | |
|---|---|---|---|
| **Assembly pieces** | **Tm** | **Size** | **Ext time** |
| Backbone | 58C | 2,812 | 90s |
| LHA | 64C | 1,902 | 50s |
| dsRed | 59C | 1,700 | 50s |
| RHA | 59C | 1,021 | 30s |

| Primers for side-directed mutagenesis (SDM) | |
|---|---|
| **Name** | **Sequence** |
| A1315V, S1318N | cttgaggaattccgagtgggaaaaaatgtaccttctgccctc |
| A1315V | ccttgaggaattccgagtgggaaaaagtgtacctt |
| S1318N | tgaggaattccgagcgggaaaaaatgtaccttctgc |
| K1335R | ggaagatgtgatattcggcaacatacgggaaatacaccacttcca |

| Sequencing primers for SDM | |
|---|---|
| Forward | TCGTCGGCAGCGTC AGATGTGTATAA AG GCAAACAGAGAGGGCG |
| Reverse | GTCTCGTGGGCTCGG AGATGTGTATAA AG TACATATCGAACTTGGAGGC |

| gRNAs for CRISPR injections | |
|---|---|
| Tr_gRNA_L1 | TTCCTTGAGGAATTTCGAGC |
| Tr_gRNA_R1 | TATAAAAACGTATTAAAGCA |

| Primers to genotype RAL386 chromosome 2 for genomic background control | |
|---|---|
| CG9932_forward | CGAACTCTGCCAAGTCACCT |
| CG9932_reverse | GCAGAAGGTATCGGATCGGG |

**Table S3. Viability of crosses between intermediate haplotypes.**

| [WT] Genotype | Cross | [WT] F1 | [Sb] F1 | Observed [WT] ratio | Expected [WT] ratio | p-value |
|---|---|---|---|---|---|---|
| ASR/ANR | ASR/Sb x ANR | 52 | 56 | 0.48 | 0.50 | 0.7892 |
| ANK/ANR | ANK/Sb x ANR | 50 | 43 | 0.54 | 0.50 | 0.6599 |
| VSK/VNK | VSK/Sb x VNK/Sb | 26 | 41 | 0.39 | 0.33 | 0.5891 |

NOTE – The viability of F1 phenotypes (WT: wild type; Sb: Stubble) were quantified for the following crosses:

ASRxANR: ♂ ASR / TM6B, P{YFP}, Sb[1] Tb[1] ca[1] x ♀ ANR / ANR

VSKxVNK: ♂ VSK / TM6B, P{YFP}, Sb[1] Tb[1] ca[1] x ♀ VNK / TM6B, P{YFP}, Sb[1] Tb[1] ca[1]

ANKxANR: ♂ ANK / TM6B, P{YFP}, Sb[1] Tb[1] ca[1] x ♀ ANR / ANR
